# Tenacibaculosis in wild-caught, captive Chinook salmon (*Oncorhynchus tshawytscha*) in British Columbia, Canada

**DOI:** 10.1101/2023.02.17.529034

**Authors:** Emiliano Di Cicco, Kaitlyn R. Zinn, Stephen D. Johnston, Karia H. Kaukinen, Shaorong Li, Jonathan F. Archambault, Kaleb N. R. Mantha-Rensi, Kayla A. J. Zielke, William S. Bugg, Gideon J. Mordecai, Arthur L. Bass, Christoph M. Deeg, Andrew W. Bateman, Scott G. Hinch, Kristina M. Miller

## Abstract

Substantial, acute mortality was observed in wild-caught Chinook salmon (*Oncorhynchus tshawytscha*) of various ages and sizes, from sub-adults to returning adults, held in tanks during two holding studies carried out in August and September 2022 at the Bamfield Marine Sciences Centre (BMSC), in Bamfield, British Columbia, Canada. Within days of capture, a substantial number of fish began presenting lethargy, loss of balance, abnormal swimming behavior, and skin ulcers involving the caudal peduncle, fins, belly, trunk and mouth. Molecular testing revealed high levels of *Tenacibaculum dicentrarchi* in skin swabs and gills, without appreciable consistent detections of other infectious agents. *T. dicentrarchi* was also isolated from the skin ulcers. Histological analysis confirmed the presence of ulcerative dermatitis and myositis, associated with mats of filamentous, rod-shaped bacteria. In two individuals, the infection became systemic, with a bacterial colony (identified as *T. dicentrarchi*) observed in the liver of one individual. In-situ hybridization against *T. dicentrarchi* and *T. maritimum* confirmed the presence of only the former agent in the gills and skin ulcers of the affected fish. This clinical report represents the first diagnosed case of tenacibaculosis in wild-caught (captive) Chinook salmon in British Columbia.

**CONFLICT OF INTEREST:** The authors have no conflicts of interest to report.

**DATA AVAILABILITY STATEMENT:** The data that support the findings of this study are available from the corresponding author upon reasonable request.

## Introduction

Tenacibaculosis is an economically important disease in wild and farmed marine fish worldwide, resulting from the bacterial infection by various *Tenacibaculum* species (Alvarez and Santos, 2018). Tenacibaculosis is a disease characterized by skin ulcers, mouth erosions and frayed fins and tail (Toranzo et al., 2005). The occurrence of tenacibaculosis in marine-reared salmonids is spread widely geographically, ranging from Norway to Chile (Avendaño-Herrera et al. 2016; Småge et al., 2016; Olsen et al., 2017; Valdes et al., 2021; Spilsberg et al., 2022), including outbreaks on open-net pen Atlantic salmon (*Salmo salar*) farm sites in British Columbia (BC), Canada (Frisch et al., 2018; CSAS-044, 2020; Bateman et al., 2021a, b; Nowlan et al., 2021a).

For Atlantic salmon in BC, tenacibaculosis often presents with yellow-pigmented plaques in the oral cavity and gills of fish, and can lead to cumulative mortality upwards of 40% in infected marine net pens (Frisch et al., 2018; Wynne et al., 2020). Some authors described this manifestation of tenacibaculosis occurring in Atlantic salmon in BC as “mouthrot” (Frisch et al., 2018; Wade and Weber, 2020; Nowlan et al., 2021b), to give more emphasis to the occurrence of yellow plaques in the oral cavity and ulcerative stomatitis which, in some individuals, appear to be the only clinical sign reported. However, in a minority of cases Atlantic salmon affected by mouthrot can also present with erosion of the skin and fins as well as ulcers (Nowlan et al., 2021b), findings that are consistent with reports of tenacibaculosis in salmonids worldwide (Valdes et al., 2021; Avendano-Herrera et al., 2006; Olsen et al., 2020; Spilsberg et al., 2022).

Early research into the causal agent of mouthrot in BC identified *Tenacibaculum maritimum* as the responsible pathogen (Frisch et al., 2018; Wynne et al., 2020), with experimental trials – using *T. maritimum* isolates collected from outbreaks on Atlantic salmon farms in BC– successfully inducing mouthrot in exposed Atlantic salmon (Frisch et al., 2018). However, new molecular assays have recently been developed to identify the presence and abundance of other *Tenacibaculum* species, including *T. dicentrarchi* and *T. finnmarkense* (Avendaño-Herrera et al. 2016; Smage et al., 2016; Bridel et al., 2018; Nowlan, 2020). These have both been detected on salmon farms, can co-occur with *T. maritimum*, and may be responsible for some cases of tenacibaculosis (Alvarez and Santos, 2018; Avendaño-Herrera et al. 2016; Smage et al., 2016, 2018; Olsen et al., 2017; Mimeault et al., 2020). On Atlantic salmon in BC, at least six species of *Tenacibaculum* have been confirmed, including *T. maritimum, T. dicentrarchi* and *T. finnmarkense* (Smage et al., 2018; Nowlan et al 2021a, b; Nowlan et al.,2023). The latter three species have been established through international challenge studies as causal agents of tenacibaculosis (Smage et al., 2018; Frisch et al., 2018; Klakegg et al., 2019; Nowlan et al., 2021b; Valdes et al., 2021). Thus, the clinical presentation of tenacibaculosis in BC could be caused by various *Tenacibaculum spp*., which in some cases may involve infection with *T. dicentrarchi* and/or *T. finnmarkense* (Nowlan et al., 2021a; Wynne et al. 2020).

There are various reports of tenacibaculosis in Pacific salmon (*Oncorhynchus spp*.), including cases from California (Chen et al., 1995), Chile (Sandoval, 2020; Valdes et al., 2021), and Alaska (Meyers et al., 2019). Further cases in Pacific salmon have been reported from BC, but these have not been well characterized (Wade and Weber, 2020). In this report we present the first diagnosis of tenacibaculosis in wild-caught Pacific salmon from BC waters. We describe the severity of disease and, through evaluation and localisation of the infectious agents present, we identify the most likely causative agent responsible.

## Material and Methods

Wild Chinook salmon were caught in Barkley Sound using recreational fishing gear for a series of holding studies to examine fishery capture and handling methods on post-release physiology and mortality. The fish analyzed in this report come from two consecutive holding studies (under DFO collection permit XR 143 2022 and UBC Animal Care approval A19-0193009). Upon capture, a blood sample was taken 15 minutes after gear hook up (for a separate study), and a subset of fish were swabbed on the flank along the lateral line, and biopsied (non-lethal gill sample). Within two hours of capture, the fish were transported to holding tanks at the Bamfield Marine Sciences Centre (BMSC; under DFO transfer permit 129330). Water in the holding tanks was pumped directly from the inlet in front of the BMSC. Surface ocean temperature at the time of capture was between 11 and 20°C, while the temperature inside the holding tanks varied between 10 and 13°C. Moreover, the tanks were disinfected with Virkon ® S (1% solution for 10min; Dynamic Aqua Supply Ltd., Surrey BC, Canada) before and between studies, to avoid contamination or cross-infection.

The fish presenting altered behavior and external lesions (detailed below) were euthanized via a blow to the head. Molecular samples were collected from all the fish involved in the two studies, while histological samples were collected from seven fish (four from the first study, and three from the second study), which were either moribund, euthanized or recently dead in the holding tanks (i.e. fresh mortalities).

Dissected gill, anterior kidney and liver tissues were collected and preserved in RNAlater for molecular testing against 47 infectious agents using high-throughput quantitative polymerase chain reaction (HT-qPCR) on the Fluidigm Biomark Dynamic ArrayTM microfluidics platform (Fluidigm, San Francisco, CA, USA) at the Pacific Biological Station, Nanaimo, British Columbia, Canada. This platform has recently been analytically validated for quantitative infectious agent profiling in salmon tissue (Miller et al. 2016) and applied to over 50 studies of Pacific salmon (e.g. Di Cicco et al. 2017; Miller et al. 2017; Thakur et al. 2018). Infectious agent taxa were chosen based on knowledge of their presence in Canada or evidence of their association with disease worldwide (Miller et al. 2016) as well as the addition of new assays for agents identified in BC salmon within the Strategic Salmon Health Initiative program (Mordecai et al. 2019). Assays utilizing Taqman probes (Table S1) were designed to target both RNA and DNA. Swabs from the edges of skin lesions were also collected and tested using the same molecular methods as the tissues.

Cryo-stocks were collected by swabbing lesions with a sterile toothpick that were placed in aged sea water with 10% glycerol, preserved by flash freezing in liquid nitrogen, and stored at -80°C. 50μl of supernatant from the glycerol stock was plated on selective marine Sheih’s Media (MSSM; Polymyxin B and Sisomicin) and selective FFM (Kanamycin) agar plates and incubated at 4°C and 20°C. Culture identity was confirmed using species-specific qPCR assays on the Quantstudio 6 platform (Applied Biosystems, MA USA); see Table S1.

Gills, anterior and posterior kidney, spleen, heart, liver and muscle sections from around the edge of skin lesions were collected, preserved in 10% neutral buffered formalin and embedded in paraffin for histological evaluation. Histology slides were prepared and stained with routine Hematoxylin & Eosin (H&E) solutions for standard pathological evaluation. Additionally, consecutive sections from the same paraffin blocks were used for in-situ hybridization (ISH).

ISH was carried out as per the manufacturer’s instructions (ACD Bio Amplification kit RNAscope 2.5HD detection reagent RED, ref. #322360) using probes against *T. maritimum* (ACD Bio RNAscope, ref. #1200621/C1) and *T. dicentrarchi* (ACD bio RNAscope, ref. #1200611/C1).

Positive and negative control probes were used as well to determine the accuracy and specificity of the test (ACD Bio RNAscope Positive CTRL Om/ppib ref. #540651; Negative CTRL DapB ref. #310043). The *T. dicentrarchi* and *T. maritimum* probes utilized for ISH included the sequences used in the species-specific qPCR assays for the respective species, as well as additional sequences unique to the respective species to ensure specificity.

## Results

### Clinical Observations

Within four days of transportation to the holding tanks, a significant portion of fish (up to 19% by the end of one of the two holding studies presented in this clinical report - see Table 1) started showing lethargy, loss of balance, abnormal swimming behavior and external lesions involving the caudal peduncle, fins, belly, trunk and mouth. These lesions were classified primarily as scale loss, skin erosions and ulcers associated with hemorrhages, and a white or yellowish margin. The fins and tails were eroded, while the ulcers involved the deeper muscle tissue (Figure 1).

**Table 1.**
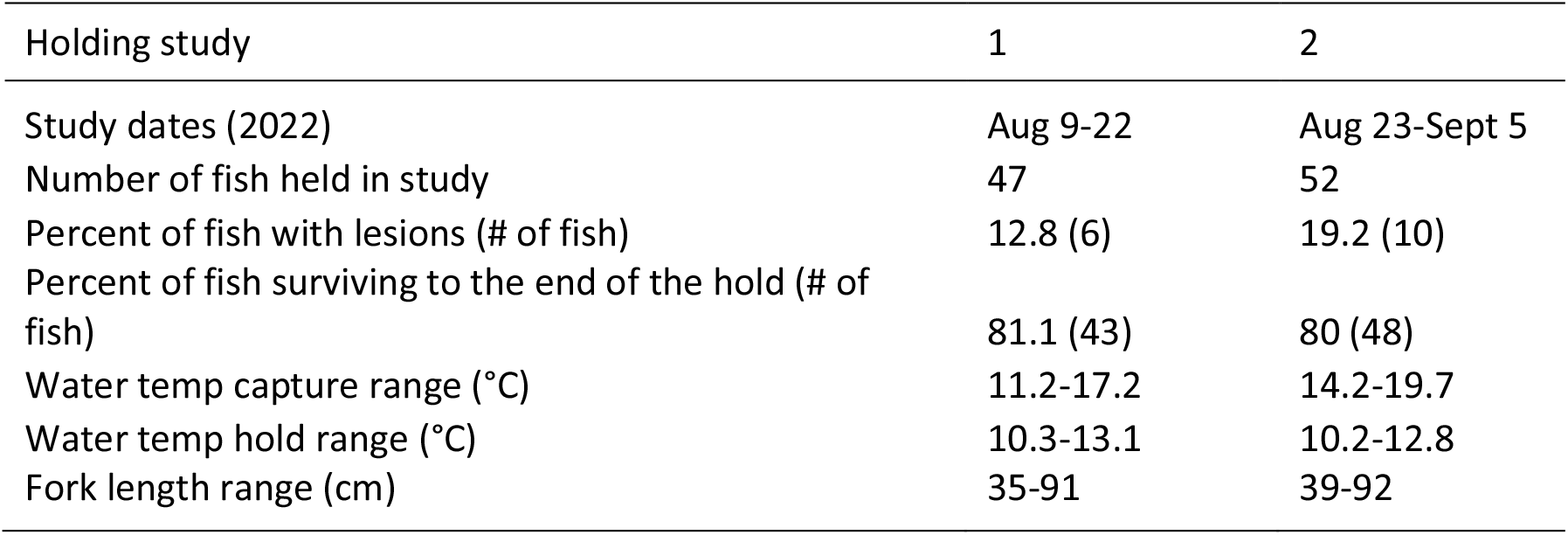
Summary of holding study parameters and Chinook salmon sampled.

**FIGURE 1.**
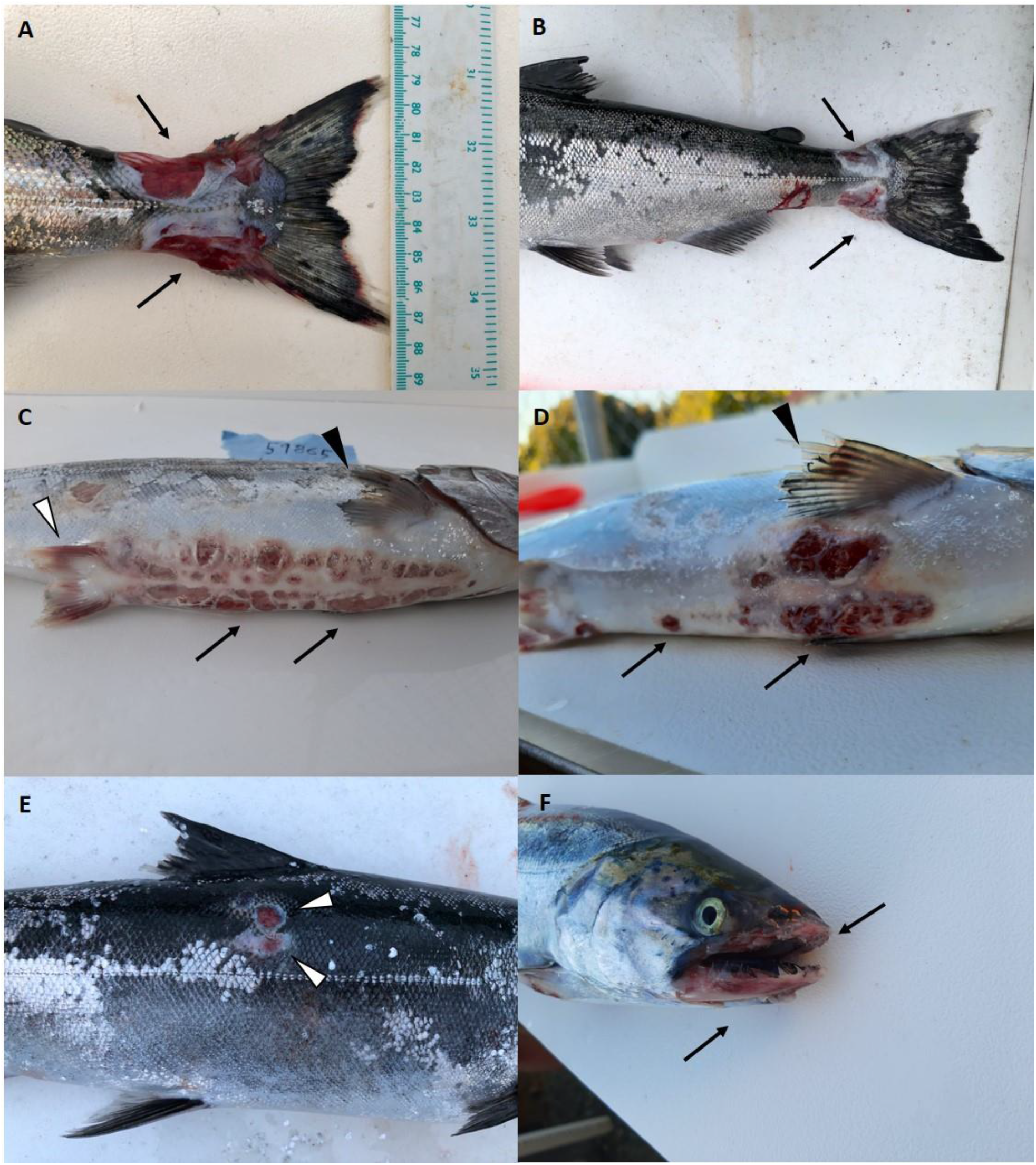
External lesions observed in the samples analyzed: A-B. Skin ulcers (A) and erosions (B) involving the caudal peduncle (arrows). C-D. Skin ulcers and erosion on the belly (arrows). Fin hemorrhages (white arrowhead – C) and erosion (black arrowheads – C, D) were also observed. E. Extensive scale loss associated with ulcerative lesions (white arrowhead) on the flank. F. Severe erosion and ulcer of the mucosa and skin on both maxilla and mandible (arrows).

### Molecular Analysis

Molecular results from the samples collected and analyzed in this study are shown in Figure 2 (results discussed here) and Table S2 (results for all fish from the two holding studies). The results we discuss in the rest of this paper refer to the seven focal fish unless indicated. Out of the 47 infectious agents tested, 18 were detected in the seven focal fish, and most only at background levels (i.e. near or below the limit of detection). Some agents, like the bacterium *Candidatus* Branchiomonas cysticola and the microsporidian parasite *Paranucleospora theridion*, were observed at high levels only in the gills at time of capture, and in the kidney during the subsequent holding study, but not in the skin swabs or gills when external lesions were observed. Alternatively, the myxozoan *Parvicapsula pseudobranchicola* was detected throughout the duration of the study, but at low levels. One agent, *T. dicentrarchi*, was not detected at the time of capture in any of the fish analyzed for histology (but was detected in three other fish across the two holding studies), while very high levels of this bacterium were observed in the skin swabs and gills of lesioned fish during holding (Figure 3). The kidneys and livers of these fish were also positive for *T. dicentrarchi*, but at a considerably lower load, with the exception of two fish where bacterial loads in the livers were similar to that in the gills and skin swabs. *T. maritimum* was detected at the time of capture in four of the seven fish which were subsequently analyzed for histology (three detections in skin swabs and three gill biopsies), but *T. maritimum* was not detected in these focal fish during holding, with the exception of one individual with a very low load detection in the gills. *T. maritimum* was, however, detected during holding in several of the cohabitant fish (Table S2). *T. finnmarkense* was not detected in any of the focal samples, but it was detected in four cohabitant fish (either at capture or during holding; Table S2). All three *Tenacibaculum* species for which we tested were detected in some fish at the time of capture (Table S2).

**FIGURE 2.**
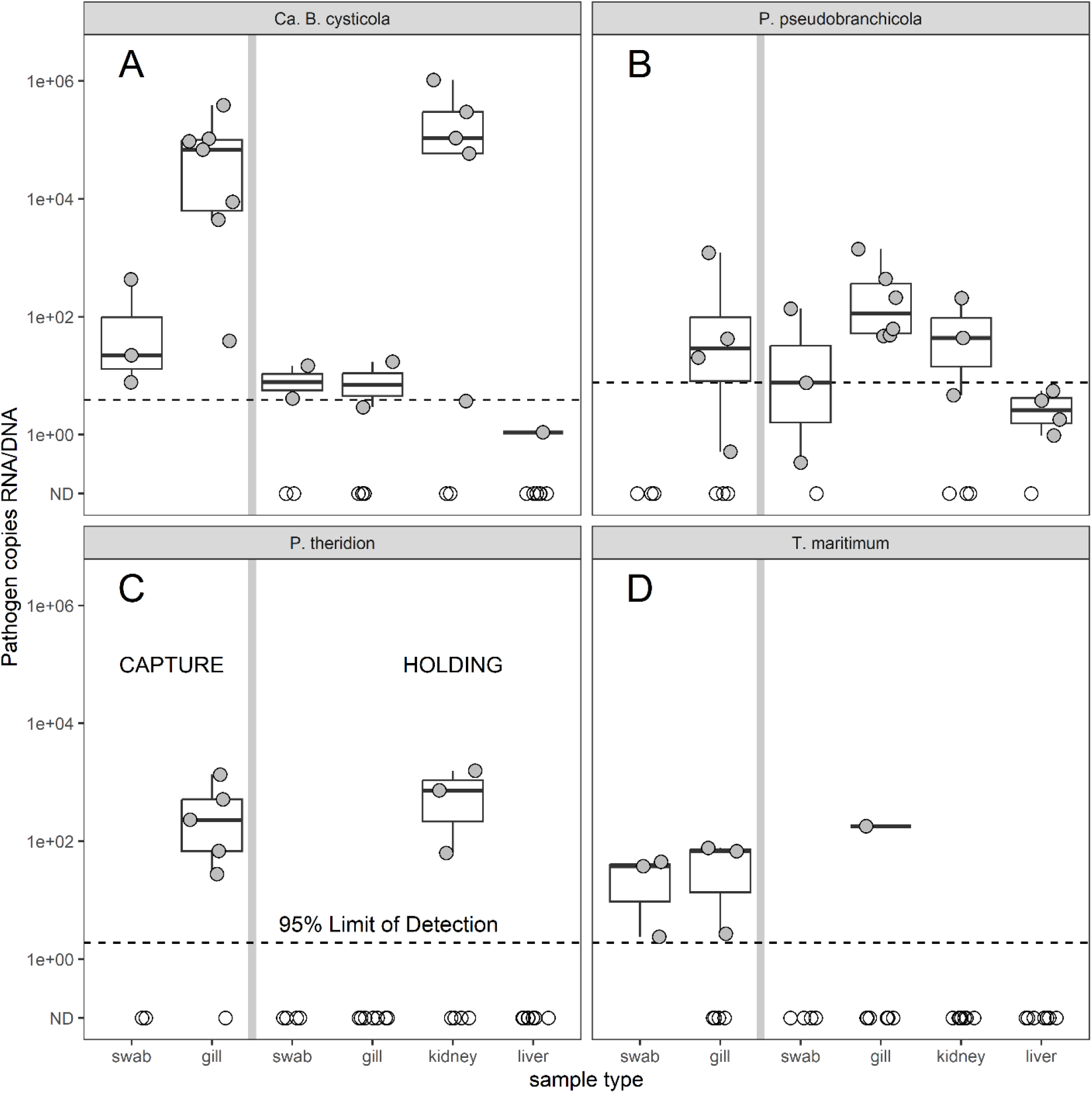
Detection of pathogenic agents via qPCR in the submitted samples. The seven focal samples tested positive for 18 infectious agents. Only five (four of which are showed in this figure) showed significant detections or trends. A.The bacterium *Candidatus* Branchiomonas cisticola was observed throughout the durantion of study, but at high levels only in the gills at time of capture, and in the kidney during the subsequent holding study. B. The myxozoan *Parvicapsula pseudobranchicola* was detected throughout the duration of the study, but at low levels. C. The microsporidian parasite *Paranucleospora theridion* was observed at high levels only in the gills at time of capture, and in the kidney during the subsequent holding study. D. *T. maritimum* was detected at the time of capture in four of the seven fish which were subsequently analyzed for histology (three detections in skin swabs and three gill biopsies), but was not detected in these focal fish during holding, with the exception of one individual with a very low load detection in the gills. All results are shown as copies per µg of RNA/DNA.

**FIGURE 3.**
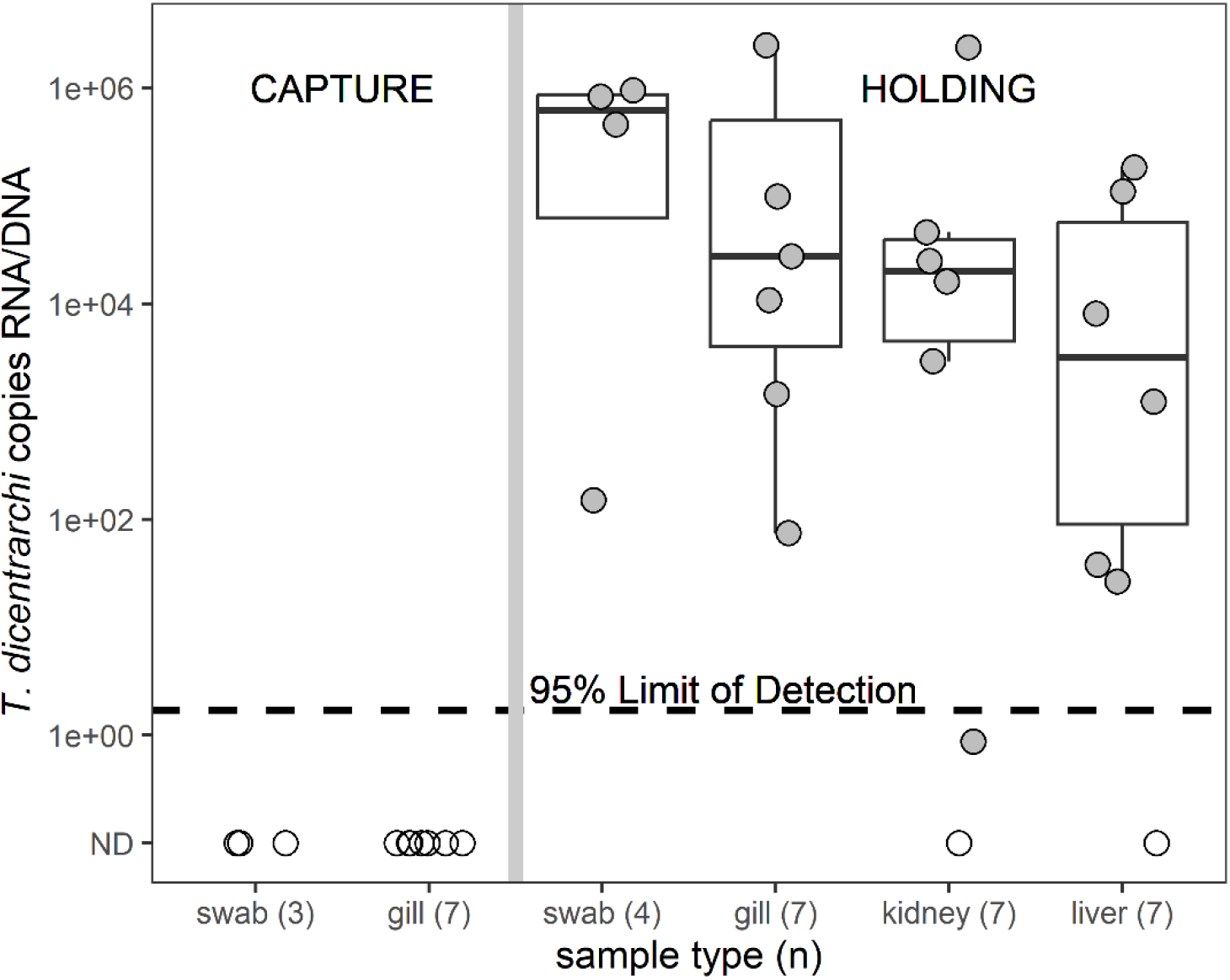
Details of the molecular detection of *Tenacibaculum dicentrarch*i in the samples analyzed. The agent was primarily detected in the gills and in the skin swabs from the wounds of the affected fish during the holding period, as well as in liver and kidney, although at a significantly lower degree. All results are shown as copies per µg of RNA/DNA.

### Bacterial isolation

Colony growth was observed on all media and temperature combinations. Upon PCR confirmation, colonies grown at 4°C on MSSM agar collected from a skin lesion on a fresh mortality tested positive for *T. dicentrarchi* (Figure S1). Colonies presented pale yellow and convex (Figure S2), consistent with observations in the literature (Klakegg et al., 2019).

### Histology

Histological analysis of the seven focal fish collected (see Doc S1 and S2 for details on single fish histological analysis, performed by the lead author and independently corroborated by a second veterinary pathologist, respectively) revealed lesions primarily affecting the skin and muscle, and consisting of ulcerative dermatitis and myositis. Overall, loss of scales, epidermis and epidermal spongiosis (when present), and deep ulcers were observed, often extending past the dermis and involving the underlying muscle. The lesions were frequently associated with mats of filamentous, rod-shaped bacteria (e.g. Figure 4A): such mats were occasionally observed infiltrating the dermis and inside the exposed muscle bundles (e.g. Figure 4C, E-F).

**FIGURE 4.**
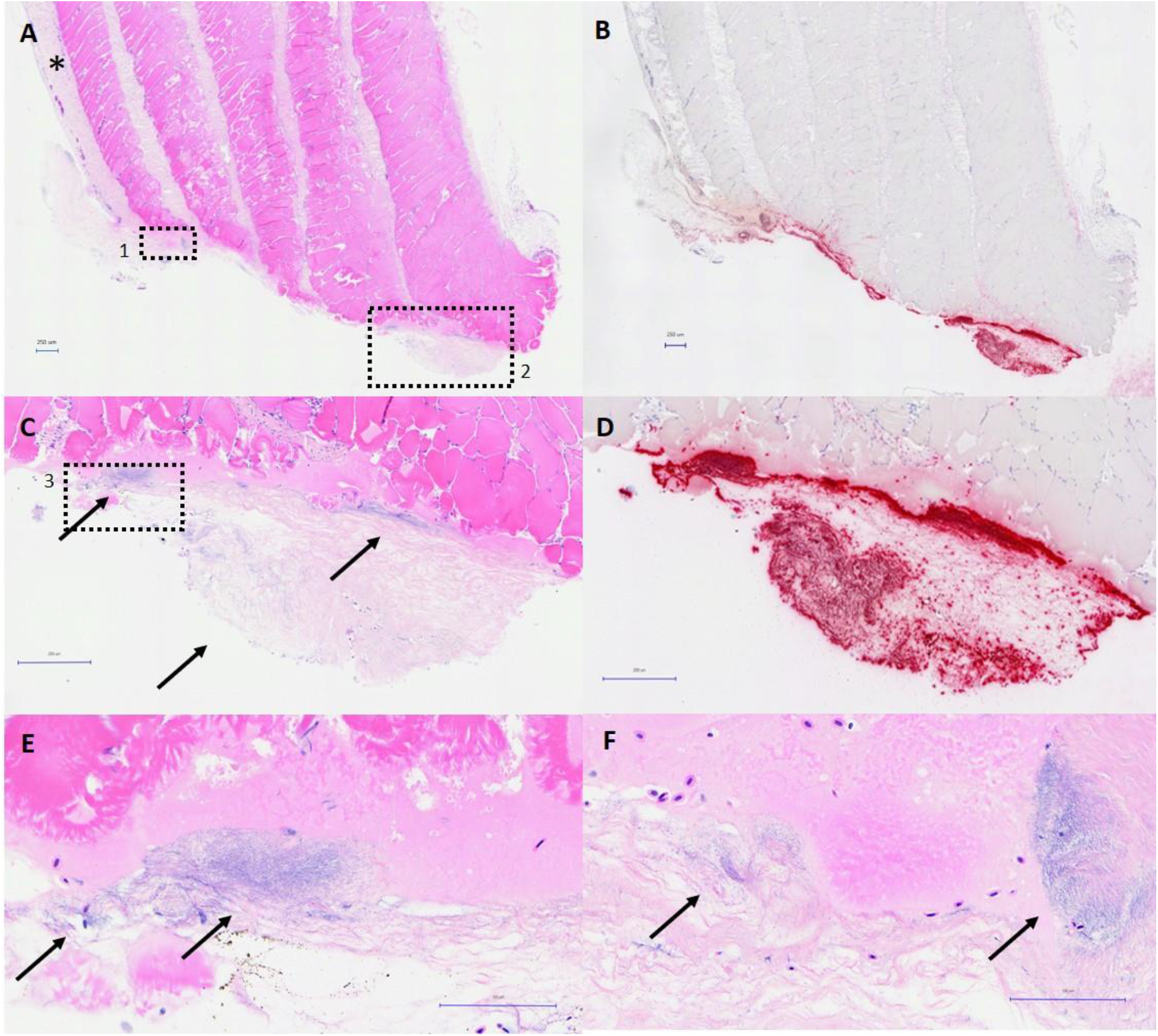
Pathology of the skin lesions (Fish ID# 59822): A. Skin ulcer with loss of epidermis and dermis, and several mats of filamentous, rod-shaped bacteria covering and infiltrating the underlying muscle bundles, some of which are degenerated. Note intact skin (*). Scale bar 250µm. B. Same field as A, marked with probe against *Tenacibaculum dicentrarchi* (ISH). Note red (positive staining) detected on the external surface of the wound, as well as a few infiltrating bacteria penetrating deeper between muscle bundles. Scale bar 250µm. C. Higher magnification of insert 2. Note the large mats of bacteria on the surface of the muscle bundles, as well as in the overlying necrotic material. Scale bar 250µm. D. Same field as C. Note red (positive staining) identifying large mats of *T. dicentrarchi*. Scale bar 250µm. E. Higher magnification of insert 3. Note the filamentous, rod-shaped bacteria replicating in the necrotic material covering degenerated muscle bundles. Scale bar 250µm. F. higher magnification of insert 1. Two large mats of filamentous bacteria in the necrotic material covering the wound. Scale bar 250µm.

The dermis (when present) displayed oedema and necrosis, while the muscle bundles involved in the infection showed myodegeneration and necrosis. The gills were also affected by the infection, with necrosis and lysis of the lamellar epithelium, filamentous bacteria infiltrating into the epithelium, and mats of bacteria proliferating in the overlaying mucus (Figure 5). There was no inflammatory response to the infection in the areas around the lesions and, in most cases, no evidence of a systemic infection. However, in one fish (fish ID 59865) a bacterial colony (identified as *T. dicentrarchi* by ISH, see below) was observed in the hepatic parenchyma (Figure 6C).

**FIGURE 5.**
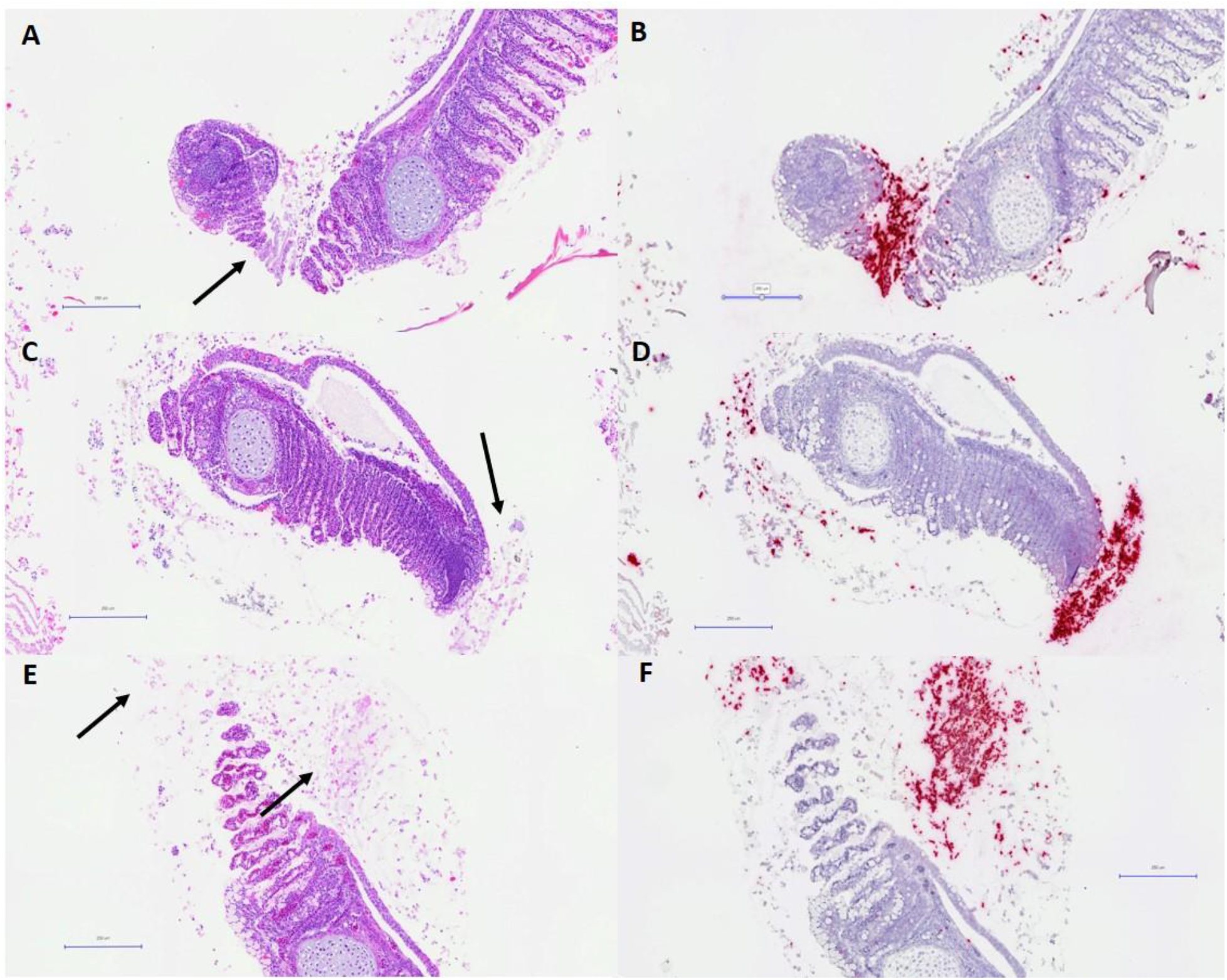
Pathology of the gills (fish ID# 59865): A, C, E. Gills showing mats of filamentous, rod-shaped bacteria in the lamellae (A), as well as in the excess of mucus covering the lamellae (C, E). H&E. Scale bar 250µm. B, D, F. Same fields as A, C, E, marked with probe against *Tenacibaculum dicentrarchi* (ISH). Note red (positive staining) detected in the areas where the bacterial mats were observed in the H&E samples. Scale bar 250µm.

**FIGURE 6.**
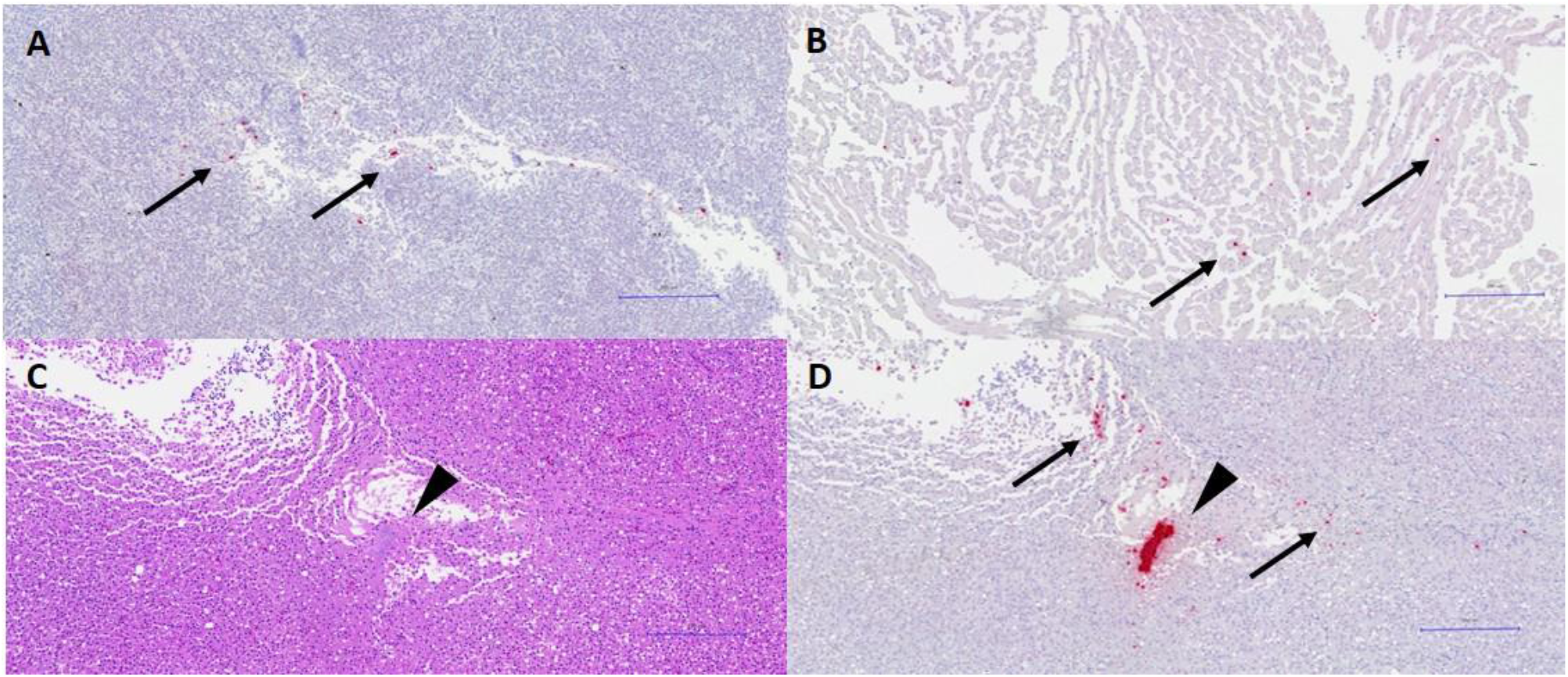
Systemic infection (fish ID# 59865): A. Section of spleen marked with probe against *Tenacibaculum dicentrarchi* (ISH). Note red (positive staining) detected where a few scattered mats of *T. dicentrarchi* was present (arrows), but not visible in the same samples stained with standard H&E. Scale bar 250µm. B. Section of heart from the same individual, showing a few clusters of *T. dicentrarchi* (arrows). Scale bar 250µm. C. Section of liver from the same individual, showing a bacterial colony localized in the parenchyma (arrowhead). No inflammatory reaction is visible. H&E. Scale bar 250µm. **(missing picture)** D. Same section as C. Note red (positive) staining detecting *T. dicentrarchi* in the bacterial colony, and other clusters of the same bacteria scattered in the parenchyma. Scale bar 250µm.

One individual (fish ID 59822) presented moderate, localized areas of parenchymal necrosis and several, widespread megalocytic hepatocytes (Figure 7). While megakaryosis was consistently present in these enlarged cells, double/multiple nuclei, pyknosis, karyorexis and sporadic intranuclear inclusion bodies were also occasionally observed.

**FIGURE 7.**
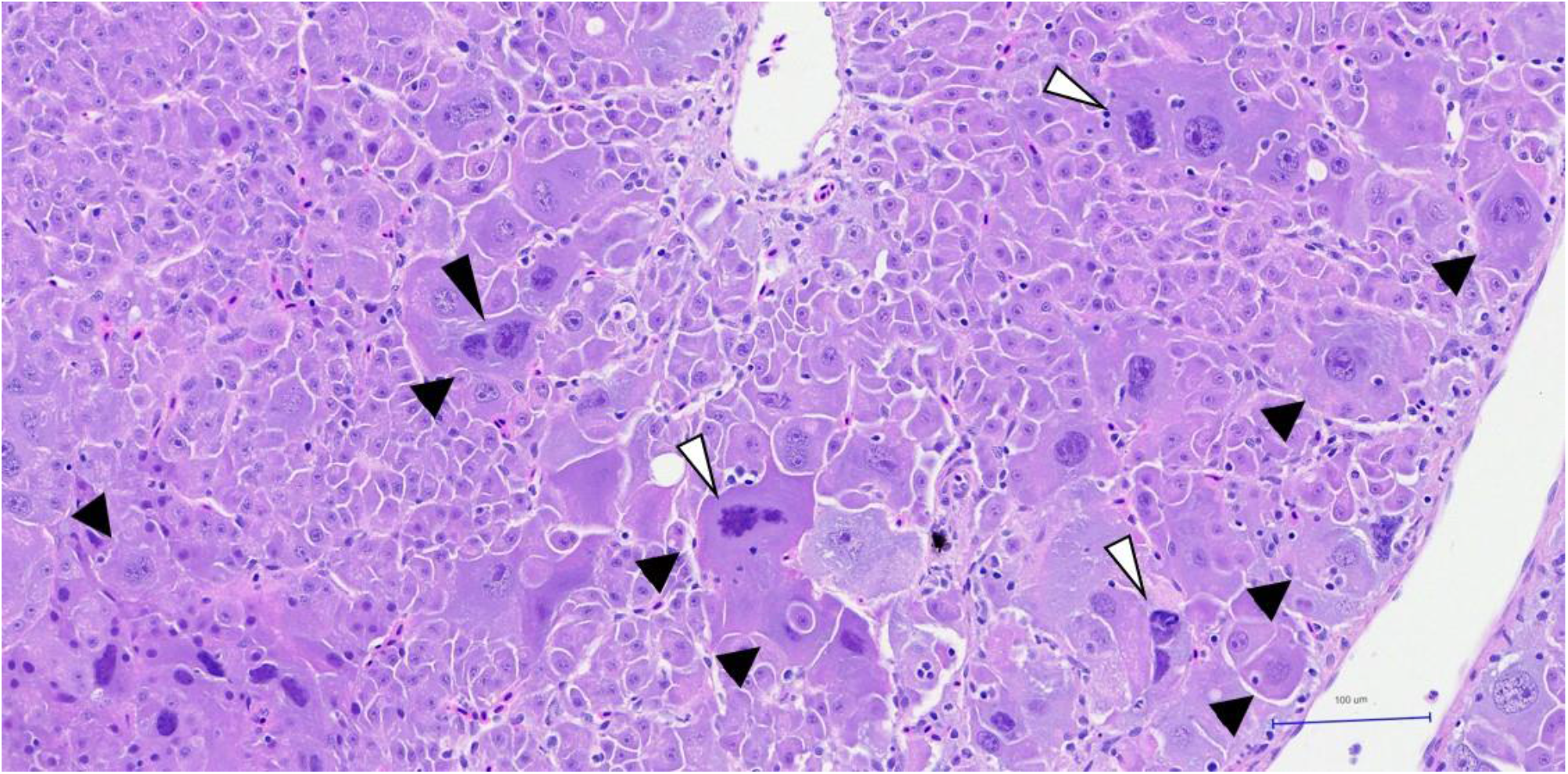
Pathology of the liver (fish ID# 59822): enlarged hepatocytes showing megalocytosis and megakaryosis scattered throughout the parenchyma (arrows). Some of these anomalous cells showed multiple nucleus (black arrowhead), while others showed pyknosis or karyorexis (white arrowheads). Scale bar 100µm.

### In situ hybridization

In situ hybridization (ISH) results are summarized in Table 2. *T. dicentrarchi* showed extensive positive staining in the skin lesions (Figure 4B). These stained areas allowed the identification of several bacterial colonies in the dermis and in the material resulting from the dermal and muscular necrosis (Figure 4D). Numerous bacteria were also observed infiltrating into the muscle bundles and between the muscle planes (Figure 4B, D). ISH also identified several bacterial colonies in the gills and associated mucus (Figure 5B, D, F). Furthermore, a bacterial colony in the liver of a single individual fish (Figure 6D) stained positive for *T. dicentrarchi*. In this same fish, further positive clusters of bacteria were found scattered in the liver as well as in the spleen and heart (Figure 6A-B), although they were not visible by examination of the H&E slides. This finding confirms the potential for *T. dicentrarchi* to induce a systemic infection.

**Table 2.** ISH results for *Tenacibaculum dicentrarchi* and *T. maritimum. T. dicentrarchi* was detected by selective ISH staining primarily in the skin and muscle of the external wounds, as well as in the gills and associated external mucus. In one individual, this agent was also found systemically (in liver, spleen and heart). There was no detection of *Tenacibaculum maritimum* in the seven focal fish undergoing pathological investigation. <u>Score legend:</u> +++: marked/abundant staining involving multiple area of the tissue; ++: moderate staining involving multiple area of the tissue; +: spotty/rare staining involving multiple area of the tissue; no score: absence of staining.

| fish ID # | *Tenacibaculum dicentrarchi* |  |  |  |  |  |  | *Tenacibaculum maritimum* |  |  |  |  |  |  |
| --- | --- | --- | --- | --- | --- | --- | --- | --- | --- | --- | --- | --- | --- | --- |
|  | Skin | Muscle | Gills | Heart | Spleen | Liver | Kidney | Skin | Muscle | Gills | Heart | Spleen | Liver | Kidney |
| 59830 | ++ | +++ | ++ |  |  |  |  |  |  |  |  |  |  |  |
| 59791 | + | ++ |  |  |  |  |  |  |  |  |  |  |  |  |
| 59822 | +++ | ++ | + |  |  |  |  |  |  |  |  |  |  |  |
| 59759 |  |  |  |  |  |  |  |  |  |  |  |  |  |  |
| 59865 | + | ++ | +++ | + | + | + |  |  |  |  |  |  |  |  |
| 59869 | n/a | +++ | ++ |  |  |  |  |  |  |  |  |  |  |  |
| 59881 | ++ | ++ | ++ |  |  |  |  |  |  |  |  |  |  |  |

As previously reported in the results for the molecular analysis, *T. maritimum* was detected in four fish used in the two holding studies analyzed in this report, as well as in several cohabitant fish during holding. For this reason, we also performed ISH against *T. maritimum* to assess whether a mixed infection may have contributed to the observed disease manifestation. *T. maritimum* was not detected by ISH in any of the samples analyzed.

### Diagnosis and discussion

The data presented in this clinical report strongly suggest that the lesions and mortality observed in wild Chinook salmon, held in tanks following capture, resulted from an infection with *T. dicentrarchi* and the consequent development of tenacibaculosis. The lesions observed in the mouth and caudal peduncle could have been initially triggered by injury associated with the removal of protective mucus during angling capture and post-capture handling procedures (i.e. hook penetration and gripping fish by the caudal peduncle). The possible interactions between disease development and different angling treatments and handling processes at the time of capture are, however, beyond the scope of this paper. Nevertheless, the observation of ulcers on skin, fins and mouth, associated with mats of filamentous, rod-shaped bacteria, and in absence of an inflammatory response (Spilsberg et al., 2022), are typical findings reported with the occurrence of tenacibaculosis (Toranzo et al., 2005). Moreover, *T. dicentrarchi* was successfully isolated from the skin ulcers. These findings add to previous cursory observations of tenacibaculosis in farmed Pacific salmon in BC (Wade and Weber, 2020), where in 2009 three farmed Chinook smolts presented with ulcerative stomatitis with abundant filamentous bacteria and yellow plaques in the mouth, during a BKD outbreak. Our findings are also consistent with characterisations of tenacibaculosis in Chinook and coho salmon (*O. kisutch*) in California (Chen et al., 1995), Chile (Sandoval, 2020; Valdes et al., 2021), and Alaska (Meyers et al., 2019).

An important feature of this case is that the lesions and mortality were most likely caused by *T. dicentrarchi*. This agent has been shown to experimentally cause disease and mortality in Atlantic salmon in BC (Nowlan et al. 2021b), and has been isolated from outbreaks of mouthrot on BC Atlantic salmon farms (Nowlan et al. 2021a). However, most studies in BC have focused their attention on the pathogenic activity of *T. maritimum* in farmed Atlantic salmon and its ability to cause mouthrot (Frisch et al., 2018). *T. maritimum* was detected in some of the focal fish from this study at the time of capture (by qPCR), but not during the holding study, when the behavioral changes and skin lesions first appeared. The negative results of our ISH against *T. maritimum* in the focal samples corroborate these findings. Despite the presence of *T. maritimum* on the skin and gills of four fish at the time of capture, and the detection of this agent in several cohabitants during holding, *T. dicentrarchi* was the only agent present in the wounds of the affected fish at remarkable levels.

It is unclear why *T. dicentrarchi* was not detected at the time of capture in the seven focal fish discussed in this report. We note that *T. dicentrarchi* was detected at the time of capture in three individuals (one in the first holding study and two in the second) that were cohabitant with the seven focal individuals. Therefore, the most likely source of infection for the focal fish could be the presence of *T. dicentrarchi* in the cohabitant fish. While there is potential for contamination of the holding tanks from previous studies, this risk was minimized by disinfecting tanks with Virkon to eliminate bacterial pathogens before each study (Lauzon et al., 2010). Importantly, the water used in the holding tanks was sourced from Bamfield Inlet, which was not pretreated before entering tanks, and therefore could have introduced new strains or species of *Tenacibaculum*. Moreover, the water in the tanks was not controlled for temperature, which could have influenced the growth of certain strains of *Tenacibaculum* over others. Preliminary evidence suggests that temperature may play a role regulating the abundance and virulence of various *Tenacibaculum* species. *T. maritimum* and *T. dicentrarchi* may be especially abundant in and around farms sites off the coast of BC in warm temperature summer months (Nowlan et al., 2021a) while *T. finnmarkense* outbreaks have been primarily associated with colder water temperatures on Norwegian farms (Spilsberg et al., 2022). Overall, diversity of the *Tenacibaculum* genus is greater than previously understood (Nowlan et al., 2023), and further genetic sequencing will be needed to understand the range of species that are involved in disease in BC and globally. Regardless of the source, or the possible impacts of environmental conditions, these findings demonstrate that *T. dicentrarchi* can result in infection, development of tenacibaculosis, and mortality in wild Chinook salmon.

One individual presented anomalous anisocytosis and anisokaryosis in a significant number of hepatocytes. Such a finding was isolated, but it could be linked to the presence of hepatotoxicants. *Tenacibaculum spp*. are a highly toxigenic group of bacteria, and in addition to the loss of osmotic integrity, which could be caused by severe skin lesions of the nature observed in this clinical report, toxemia is also considered to be responsible for many mortalities with these infections (Baxa et al., 1988; Van Gelderen et al., 2009). However, no previous controlled studies with any *Tenacibaculum* species has ever reported such an unusual finding. Ultimately, we cannot identify the source of putative hepatotoxins, which we hypothesize could be either anthropogenic (Myers et al., 1987) or naturally occurring (e.g. algatoxins such as microcystin; Kent 1990; Andersen et al., 1993). Alternatively, the presence of sporadic intranuclear inclusion bodies could support a hypothesis of a potential infection with an unknown viral agent, undetected in the infectious agent screening. Further tests will be required to assess the cause of these lesions.

## Conclusions

Taken together, our results present a clear picture. Molecular testing of skin swab, gill, liver and kidney samples from wild-caught Chinook salmon identified multiple infectious agents. However, only one agent — *T. dicentrarchi* — was consistently present at substantial loads in fish during holding periods, in tissues affected by acute ulcers, in liver and kidney samples of two fish (confirming a systemic infection in those individuals), and was isolated from skin ulcers. The histopathological analysis of skin lesions revealed the presence of ulcerative dermatitis and myositis (in absence of inflammatory reaction) associated with mats of filamentous, rod-shaped bacteria — typical features of tenacibaculosis. ISH confirmed the abundant presence of *T. dicentrarchi* in the skin ulcers and in the gills of infected fish. The overwhelming evidence points to *T. dicentrarchi* as the causative agent of these consistently observed clinical signs.

As far as we are aware, these findings provide the first diagnosis of tenacibaculosis and first observation of *T. dicentrarchi* in wild-caught Pacific salmon in BC.

## Supporting information

Doc S1

Doc S2

## Supplementary material

**Table S1**: Table containing the main information regarding the assays used to test for the 47 infectious agents (pathogens): pathogen abbreviation, pathogen name, pathogen type, accession number, primer forward sequence, primer reverse sequence, probe, assay reference.

**Table S2**: Molecular results against the 47 pathogens for all the samples collected during the two holding studies. Grey cells identify the results for the seven focal samples undergoing histological evaluation and ISH.

**Doc 1**: Detailed results of the histological evaluation of the seven focal fish (lead author)

**Doc2**: Detailed results of the histological evaluation of the seven focal fish (independent veterinary pathologist)

**FIGURE S1**: Particular of the sample location of successful *T. dicentrarchi* isolation. Cryosamples were collected by swabbing a lesion between the pectoral fins of a fresh mortality.

**FIGURE S2**: Appearance of *T. dicentrarchi* colony isolated on MSSM medium from the lesion showed in Figure S1.

## Notes

### Competing Interest Statement

The authors have declared no competing interest.

