## Supplementary material for "Tenacibaculosis in wild-caught, captive Chinook salmon (*Oncorhynchus tshawytscha*) in British Columbia, Canada": Doc S1

**Histological Analysis – Dr. Emiliano Di Cicco**

**Fish ID: 59830**

Gills: diffused lysis of lamellar epithelium; a few aggregates of filamentous bacteria on the tips of some secondary lamellae

Skeletal muscle/skin: Large, deep epidermal ulcers, with spongiosis and dermis oedema; mats of intralesional bacteria covering and penetrating the exposed dermis and underlying musculature

Heart: a few foci of hypereosinophilic myocardial cells indicating loss of striation.

Liver: NAD

Spleen: mild congestion

Kidney: N/A

**Fish ID: 59791**

Gills: NAD

Skeletal muscle/skin: small epidermal ulcer, with spongiosis and dermis oedema; mild myonecrosis; mats of intralesional bacteria covering and penetrating the exposed dermis and underlying musculature

Heart: a few foci of hypereosinophilic myocardial cells indicating loss of striation.

Liver: mild cell individualization and apoptosis; scattered areas and single cells with hypereosinophilic cytoplasm and moderate parenchymal necrosis

Spleen: mild congestion; mild white pulp hyperplasia

Kidney: NAD

**Fish ID: 59822**

Gills: normal (post-mortem detachment of the lamellar epithelium)

Skeletal muscle/skin: Large, deep ulcer with severe myonecrosis and hemorrhages; large vesicles and spongiosis in the epidermis (when present); dermis oedema and necrosis (when present); mats of intralesional bacteria covering and penetrating the exposed dermis, hypodermis and underlying musculature; (Fig. 4A, C, E-F)

Heart: two small foci lympho/histiocytic inflammation; a few foci of hypereosinophilic myocardial cells indicating loss of striation.

Liver: a few individualized hepatocytes; several, widespread enlarged hepatocytes presenting megalocytosis and megakaryosis, at times with double/multiple nucleus, pyknosis or karyorexis; moderate, localized areas of parenchymal necrosis; sporadic intranuclear inclusion bodies (Fig. 7)

Spleen: moderate, diffused congestion; mild ellipsoid necrosis, increased deposition of hemosiderin

Kidney: diffused, moderate hemopoietic tissue necrosis; local mild hemopoietic tissue hyperplasia

**Fish ID: 59759**

Gills: NAD

Skeletal muscle/skin: NAD

Heart: a few, small foci of sub-endocardial lymphocytic infiltration, at time associated with loss of striation

Liver: a few individualized, pre-necrotic hepatocytes; local areas of parenchymal necrosis

Spleen: moderate, diffused hyperplasia of the white pulp

Kidney: moderate, diffused hyperplasia of the hemopoietic tissue

**Fish ID: 59865**

Gills: severe, widespread lysis and necrosis of the lamellar epithelium; multiple mats of filamentous bacterial in the mucous, as well as (locally) in the lamellar epithelium (Fig. 5A, C, E)

Skeletal muscle/skin: loss of epidermis and dermis; oedema in the hypodermis; several bacterial colonies in the hypodermis and infiltrated in the underlying musculature; mild myodegeneration

Heart: NAD

Liver: widespread cell individualization and apoptosis (likely post-mortem); moderate diffused parenchymal necrosis and vacuolar degeneration; one small, distinct bacterial colony inside the parenchyma (Fig. 6C)

Spleen: moderate congestion

Kidney: diffused, moderate hemopoietic tissue necrosis; diffused, moderate renal tubule degeneration and necrosis; mild congestion

**Fish ID: 59869**

Gills: NAD; a few small mats of filamentous bacteria in the mucous

Skeletal muscle/skin: total loss of epidermis and dermis; several mats of filamentous bacteria infiltrating the musculature; myodegeneration and necrosis

Heart: NAD

Liver: a few individualized hepatocytes; local areas of parenchymal necrosis

Spleen: mild, diffuse congestion; mild localized hyperplasia of the white pulp

Kidney: moderate diffused lymphocytic nephritis; local hemopoietic tissue necrosis

**Fish ID: 59881**

Gills: severe, diffused lysis of the lamellar epithelium, often leading to total loss of the structure of the secondary lamellae: small, diffused mats of filamentous bacteria in the mucous and, occasionally, in the lamellar epithelium

Skeletal muscle/skin: diffused loss of epidermis and spongiosis; diffused degenerated/necrotic dermis with oedema, often associated with mats of bacteria infiltrating the layer and the underlying musculature; myodegeneration and necrosis; metazoan organism encysted in the musculature, surrounded by a mild fibrotic reaction and nearby moderate myodegeneration

Heart: NAD

Liver: severe, diffused necrosis and cell individualization

Spleen: mild stromal hyperplasia

Kidney: moderate widespread congestion; moderate, diffused renal tubules vacuolar degeneration and necrosis; mild lymphocytic infiltration; mild hemopoietic tissue hyperplasia

*NAD: no detected anomalies
