## Supplementary material for "Tenacibaculosis in wild-caught, captive Chinook salmon (*Oncorhynchus tshawytscha*) in British Columbia, Canada": Doc S2

**_Dr. Hugh W. Ferguson_**
Balquhern, Dollar
FK14 7PT, Scotland, UK

*Your Case no*: Various
*My case no*: 013123-1-11
*Date*: Jan 31st, 2023

***What?*** Selected samples of chinook salmon forwarded for second opinion from Dr. E. DiCicco, Canada. The fish selected included 310, 881, 865 & 869 (2 slides each), while the remaining 3 fish had only one slide each (311, 312 & 319). Each set of 11 slides underwent H&E staining, as well as ISH for *T. mar* and *T. dic*.

***History:*** Significant, acute mortality was observed in Chinook salmon (of various size, from sub-adults to pre-adult and returning adult) held in holding tanks at the Bamfield Marine Sciences Centre (BMSC). Fish were caught in the nearby Barkley sound using recreational gear, and within 2h from capture were transported to the holding tanks. Surface ocean temperature at capture was about 12C. The water temperature in the holding tanks was 11-13C. The water in the holding tanks was pumped directly from the inlet in front of the BMSC. Within 3-4 days from transportation to the holding tanks, a significant portion of fish (up to 10% by the end of the holding study) started showing lethargy, loss of balance, abnormal swimming behaviour, and external lesions involving the caudal peduncle, fins, belly, trunk and mouth. Such lesions were classified primarily as scale loss, erosions and ulcers, associated with hemorrhages, and a white or yellowish edge around the wounds. Ulcers around the caudal peduncle were the predominant lesions observed. The fins were often eroded down to the fin rays, while the ulcers involved the deeper muscle tissue. The fish presenting altered behaviour and lesions were euthanized, while some individuals died "naturally" in the holding tanks.
*Samples collected and analysis performed*: Fish presenting the external lesions were euthanized by a blow to the head, and samples were collected form both moribund, euthanized fish as well as fish who recently died in the holding tanks (*i.e.* fresh mortalities). Dissected gills, anterior kidney and muscle sections from around the edge of skin lesions were collected and preserved in RNAlater for molecular testing with the DFO Molecular Genetics Lab panel of infectious agents (34 agents, screened via quantitative polymerase chain reaction [qPCR]; Miller *et al*. 2016). Gills, anterior and posterior kidney, spleen, heart, liver and muscle sections from around the edge of skin lesions were collected and preserved in formalin for histological evaluation. Histology slides were prepared and stained with routine Hematoxylin & Eosin (H&E) solutions for standard pathological evaluation, while consecutive sections of the same slides were used for In situ hybridization (ISH) with probes against *Tenacibaculum maritimum* and *Tenacibaculum dicentrarchi*. Swabs from the edges of the areas showing skin lesions were also collected for molecular testing of infectious agents present in the wounds.

***Histopathology***: Most fish had severe ulcerative and necrotizing dermatitis and myositis, with bacteria present within the lesions, typical of *Tenacibaculum* spp., and correlating well with the gross observations. Most of these lesions stained well for *Tenacibaculum dicentrarchi*, and not *T. maritimum*. In addition to these lesions, one fish (312) had a most unusual cytomegalic hepatopathy. In detail:

***Dr. Hugh W. Ferguson***
Balquhern, Dollar
FK14 7PT, Scotland, UK

*Fish 310*: severe necrotic myositis with bacteria tracking down the fascial planes. These lesions stained positively for *Tenacibaculum dicentrarchi* but not *T. maritimum*. Gills were congested while other tissues had no significant abnormalities (NSAD).
*Fish 881*: Similar to 310, but not as well preserved (autolysis).
*Fish 865*: Similar to 310. Again, autolysis was an issue.
*Fish 869*: Similar to 310, but with the addition of an encysted digenean metacercaria in one fascial plane. Once again, the lesions were *Tenacibaculum dicentrarchi* positive, and *T. maritimum* negative.
*Fish 311*: no significant abnormalities detected in any tissue.
*Fish 312*: Similar to 310. In addition to the necrotic myositis however, there was a widespread and most unusual hepatopathy, comprising marked anisocytosis, with pronounced cytomegaly and even early-stage syncytial formation. There was no accompanying inflammation.
*Fish 319*: No significant abnormalities.

***Diagnosis***: In most fish, severe necrotizing bacterial myositis, entirely consistent with *Tenacibaculum* spp. One fish (312) cytomegalic hepatopathy.

***Comment***: These skin and muscle lesions are entirely compatible with *Tenacibaculum* spp. and it was interesting to see that it was *T. dicentrarchi* rather than *T. maritimum* that dominated the findings. As you are aware, *Tenacibaculum* are a highly toxigenic group of bacteria, and besides the loss of osmotic integrity that these sorts of severe skin lesions cause, toxaemia is also considered to be responsible for many mortalities with these infections. The single encysted digenean metacercaria (fish 869) is incidental. There is no doubt in my mind that these lesions are severe enough to be considered the cause of mortality; moreover there were no other lesions suggesting an underlying problem, except possibly in fish 312.

The significance of the liver lesions in fish 312 is harder to assess, although I doubt if they had any major bearing on the cause of mortality in this fish. The lack of any significant accompanying inflammation tends to mitigate against there being a viral or indeed a toxic aetiology. Nevertheless, I cannot easily explain such a pronounced response as being due to bacterial toxins (not normally seen) nor as a repair/regeneration response (again, not normally seen even when there has been severe liver damage). Microcystins from water-borne algae are another possibility – sometimes described as “net-pen liver disease”.  At this stage, all options are open.

Dr. Hugh W. Ferguson
Professor of Veterinary Pathology

*Dr. Hugh W. Ferguson*
Balquhern, Dollar
FK14 7PT, Scotland, UK

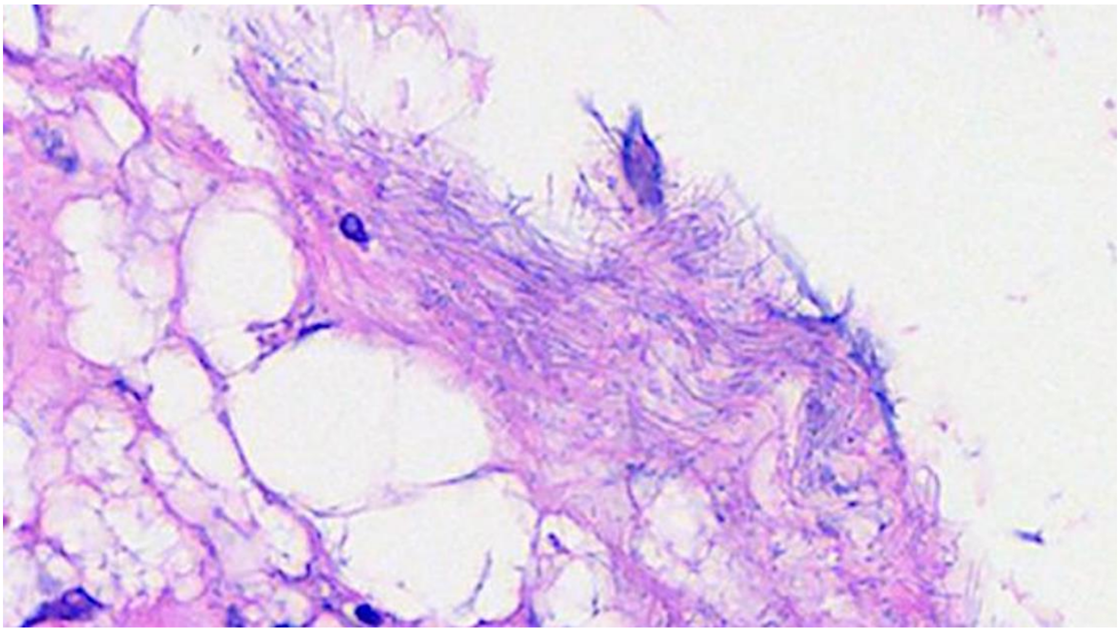

***Figure 1.*** **Typical appearance of filamentous bacteria associated with the muscle lesions.**

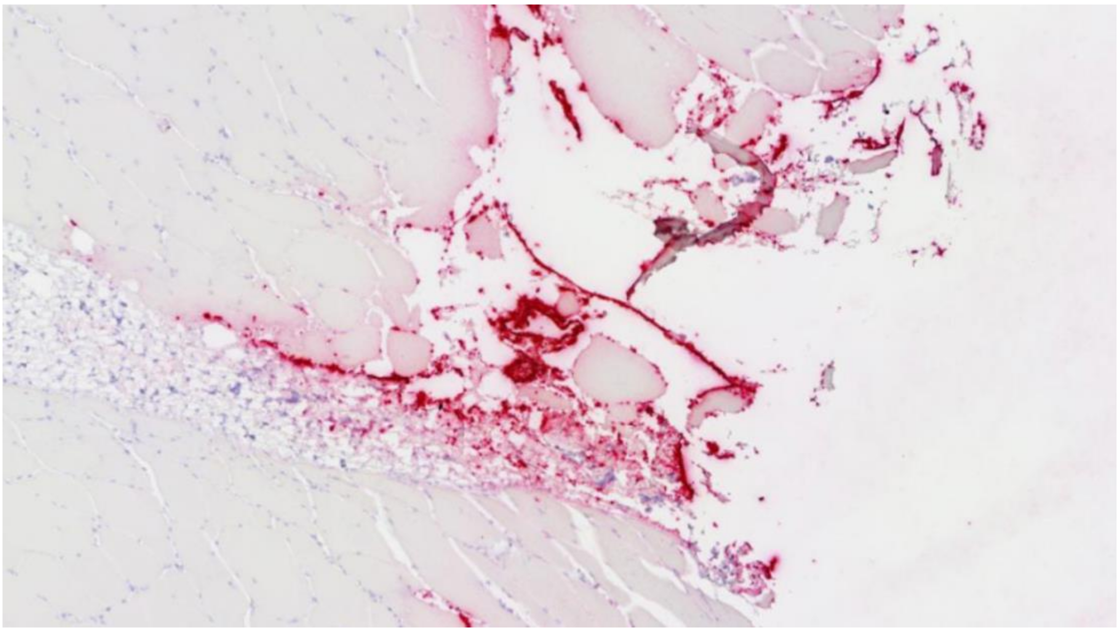

***Figure 2.*** **Lesion staining positively for** *Tenacibaculum dicentrarchi (red).*

**_Dr. Hugh W. Ferguson_**
Balquhern, Dollar
FK14 7PT, Scotland, UK

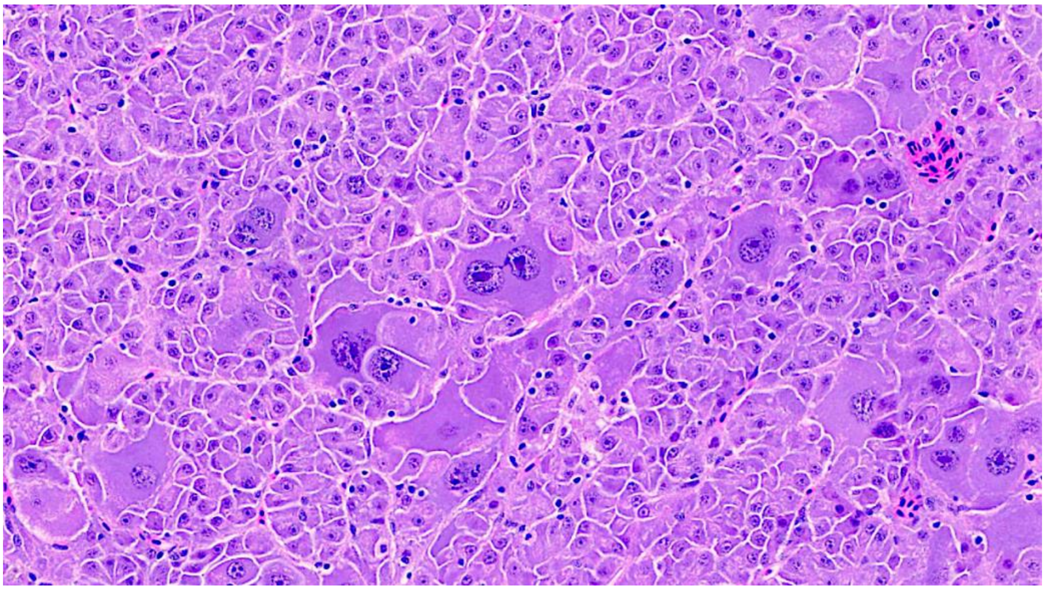

***Figure 3.*** **Marked anisocytosis in liver of fish 312.**

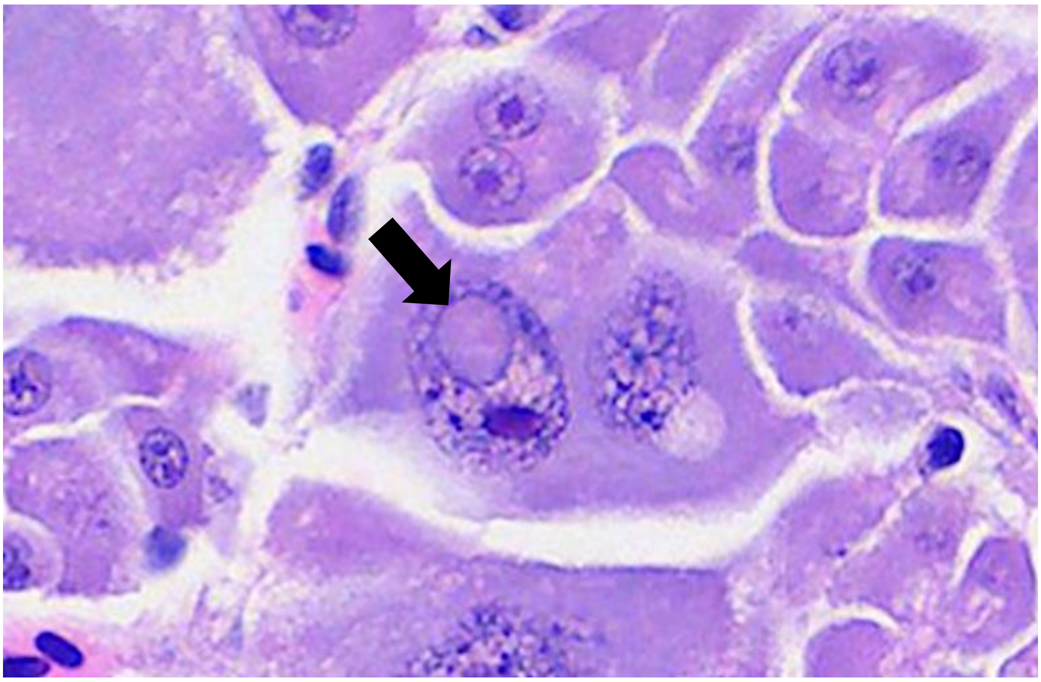

***Figure 4.*** **Well-circumscribed intra-nuclear inclusion (arrow). Virus-like or toxin-induced?**
