## Supplementary figures and images for "Tenacibaculosis in wild-caught, captive Chinook salmon (*Oncorhynchus tshawytscha*) in British Columbia, Canada"

### Fig S1

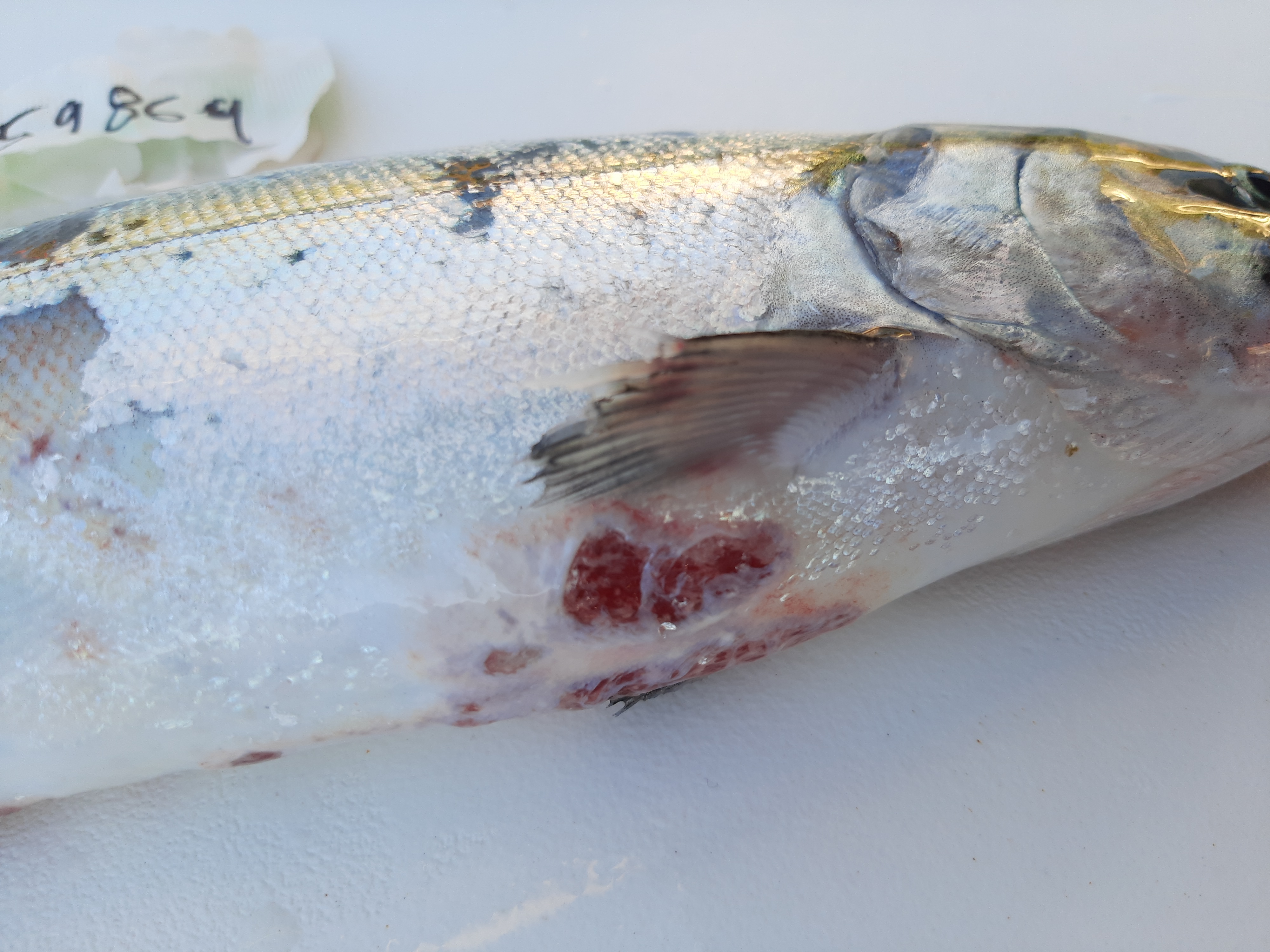

### Fig S2

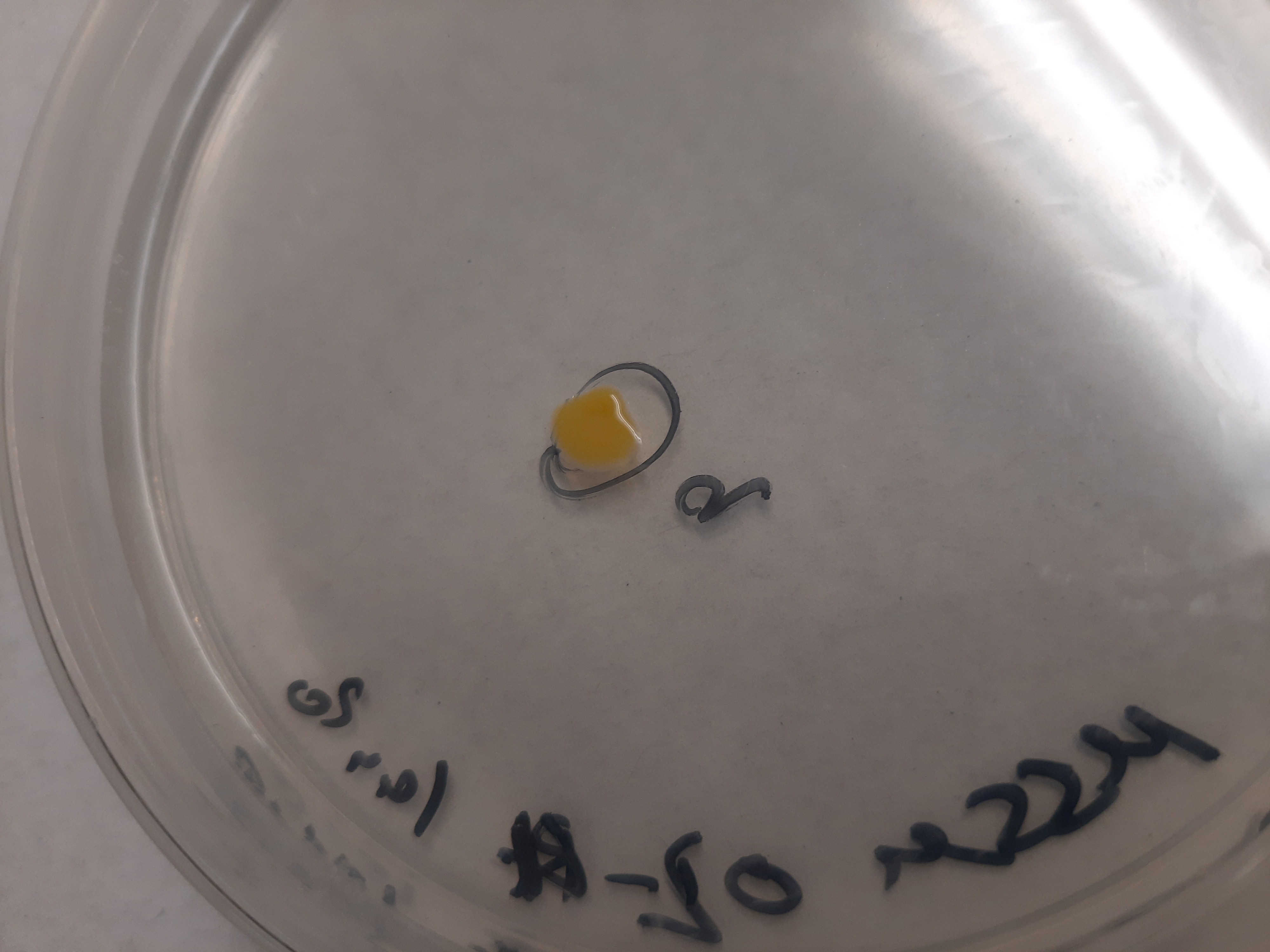
